# Printing, Characterising, and Assessing Transparent 3D Printed Lenses for Optical Imaging

**DOI:** 10.1101/2023.11.22.568002

**Authors:** Liam M. Rooney, Jay Christopher, Ben Watson, Yash Susir Kumar, Laura Copeland, Lewis D. Walker, Shannan Foylan, William B. Amos, Ralf Bauer, Gail McConnell

**Affiliations:** Strathclyde Institute for Pharmacy and Biomedical Sciences, University of Strathclyde, Glasgow, G4 0RE, UK; Department of Electronic and Electrical Engineering, University of Strathclyde, Glasgow, G1 1XW, UK; Department of Physics, University of Strathclyde, Glasgow, G4 0NG, UK

**Keywords:** Open Microscopy, Additive Manufacturing, Optics

## Abstract

High-quality lens production has involved subtractive manufacturing methods for centuries. These methods demand specialist equipment and expertise that often render custom high-grade glass optics inaccessible. We aimed to develop a low-cost, accessible, and reproducible method to manufacture high-quality three-dimensional (3D) printed lenses using consumer-grade technology. Various planoconvex lenses were produced using a consumer-grade 3D printer and low-cost spin coating setup, and printed lenses were compared to commercial glass counterparts. A range of mechanical and optical methods are introduced to determine the surface quality and curvature of 3D printed lenses. Amongst others, high-resolution interference reflection microscopy methods were used to reconstruct the convex surface of printed lenses and quantify their radius of curvature. The optical throughput and performance of 3D printed lenses were assessed using optical transmissivity measurements and classical beam characterisation methods. We determined that 3D printed lenses had comparable curvature and performance to commercial glass lenses. Finally, we demonstrated the application of 3D printed lenses for brightfield transmission microscopy, resolving sub-cellular structures over a 2.3 mm field-of-view. The high reproducibility and comparable performance of 3D printed lenses present great opportunities for additive manufacturing of bespoke optics for low-cost rapid prototyping and improved accessibility to high-quality optics in low-resource settings.

## Introduction

Additive manufacturing, in particular 3D printing, has resulted in a range of innovative open hardware solutions for optical imaging applications. The variety of printing modalities, commitment to open sharing by users, low barrier-to-entry, and rapid evolution of 3D printing technologies has provided new opportunities to make imaging more accessible, particularly in low-resource settings [1–4]. Open microscopy initiatives have aimed to provide improved access to 3D printed microscopy hardware for use in the field or in rapid clinical diagnostics, such as the OpenFlexure project [5–8]. However, such initiatives routinely focus on 3D printing the mechanical parts of the microscope, such as the chassis, focusing assemblies, or specimen mounts, mainly using fused deposition modelling (FDM) printing to manufacture parts from heated plastic filaments. The optical elements in these applications still use glass objectives or plastic camera lenses. There is a need for accessible and robust methods to produce high-quality optical elements.

Optical imaging has relied on the use of ground glass lenses to manipulate light for centuries[9–12]. Manufacturing such glass lenses is a subtractive process that can be prohibitively expensive for custom optics, requires specialist equipment and expertise, and produces a product that is delicate and easily damaged. Additive manufacturing, specifically 3D printing, has the potential to mitigate each of these barriers. Recently, additive manufacturing methods using injection moulding, magnetorheological, and molten glass printing have successfully produced bespoke optical elements [13–16]. In the latter, glass filaments are heated and extruded as in FDM printing, annealed, and cured in a kiln to produce the final product. However, this process is prohibitively costly owing to the specialist equipment and expertise required to manipulate glass in such ways. Resin-based printing techniques provide a viable solution to create bespoke optical elements for use in open hardware and imaging applications.

A variety of resin-based printing methods are available, mainly based on stereolithography (SLA) techniques. Two-photon polymerisation is routine for microfabrication of lenses, particularly for microlens arrays and x-ray imaging optics [17–19]. However, the requirement for specialist ultrashort pulsed laser sources and expensive printing instrumentation serves as a barrier to entry for these methods. Masked SLA (MSLA) printing is one of the most accessible SLA techniques [20]. This method uses a proprietary mix of methacrylate-based resin that is available in transparent and opaque forms and photoinitiated cross-linkers to polymerise under irradiation using 405 nm light. The illumination pattern is provided by an array of UV light emitting diodes, which can be collimated to provide homogeneous illumination over the entire build area. A liquid-cyrstal display (LCD) screen then masks the structure of discrete layers of the print design, changing in unison with the axial position of a buildplate mounted to a motorised stage to create a printed 3D structure [21,22]. Recent studies have produced MSLA 3D printed lenses for spectrophotometry applications [23] and described methods to quantify their material properties [24]; however, the optical performance of 3D printed lenses remains uncharacterised. Current advances in consumer-grade MSLA printer technology enable printing with a lateral resolution of up to 18 µm and an axial resolution of up to 10 µm.

We aimed to create a robust method to manufacture and characterise the optical performance of transparent 3D printed high-quality bulk optics using a consumer-grade printer and commercially available resin. We used a low-cost MSLA 3D printer and employed a spin-coating method render the lenses transparent by minimising surface imperfections and reduce scattering and refraction from step structures originating from the printing process (the so-called ‘staircase effect’). We printed a range of planoconvex lenses of various optical prescriptions, we characterised their surface profile in comparison to their commercial glass counterparts, and we assessed their optical performance.

## Methods

### 3D printing

All 3D printing was conducted using a consumer-grade Mars 3 printer (Elegoo, China) fitted with a magnetic build plate attachment for easy print removal (Sovol, China). Clear resin (Clear Resin V4; Formlabs, USA) was used as the substrate for all prints. The print settings were optimised using the Cones of Calibration test print (TableFlip Foundry) and the print quality was verified using the Ameralabs Town test print (MB Labsamera, Lithuania) (Supplementary Figure 1). All lens prints were conducted using the parameters detailed in Supplementary Information, with the planar side of the lenses printed directly on the buildplate surface. Figure 1 provides a graphical overview of the manufacturing process for 3D printed lenses.

**Figure 1.**
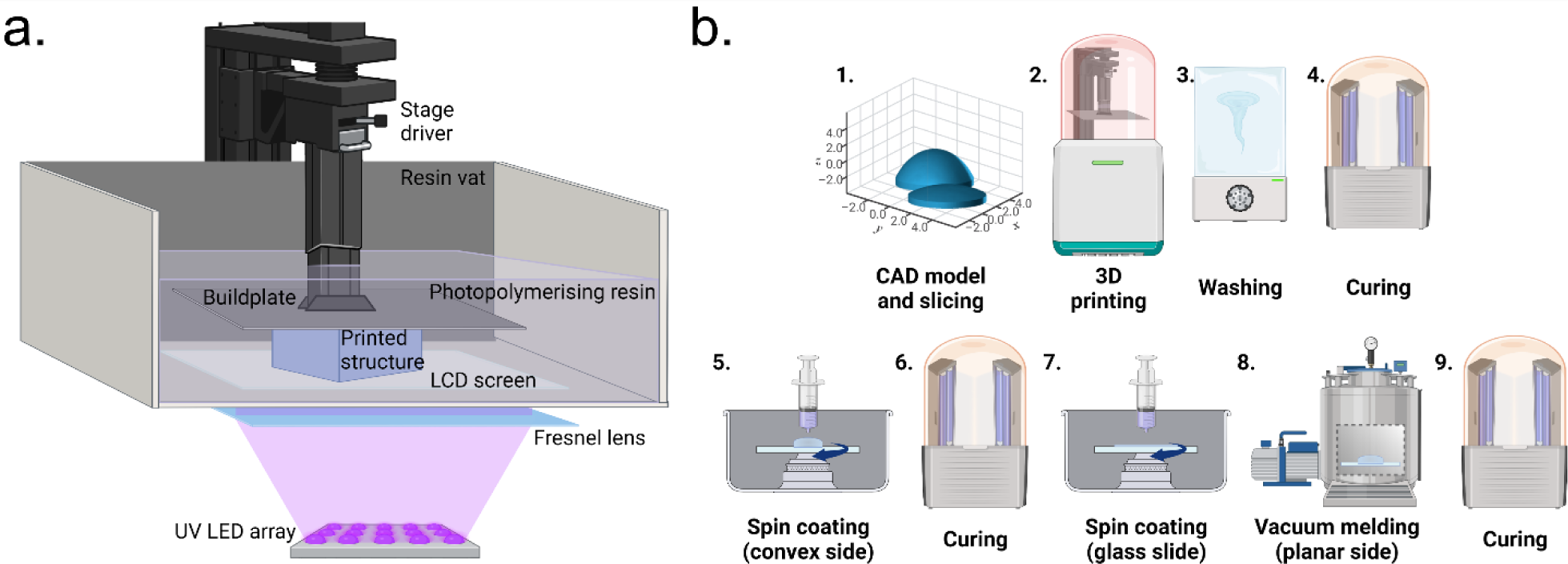
Methods for Manufacturing 3D Printed Lenses. **(a)** A schematic of an MSLA printer. A collimated UV light source is projected onto an LCD screen which illuminates and masks individual print layers as the buildplate is lifted from a vat of photopolymerising resin. **(b)** A schematic of manufacturing process for 3D printed lenses. The initial design and printing process is conducted (1-4) before a series of spin coating and curing steps to provide a smooth lens surface and increased optical quality (5-9).

Planoconvex lens designs (each with a lens diameter of 12.5 mm) were acquired from the manufacturer (Thorlabs, USA or Edmund Optics, USA) as .STEP files. Files were imported into Fusion 360 (v2.0.16985; Autodesk, USA) and the polygon count was increased to the maximum available before being exported as .STL files. Files were then imported into LycheeSlicer (v5.2.201; Mango3D, France), where the print parameters were applied, and the print files were exported as .CTB files. The .CTB lens files were printed with a raft surrounding the lens perimeter which provided a handling surface to avoid touching the lens surface. This increased the total diameter of the print to 25 mm, and ensured the print would fit in a standard 25 mm diameter optical mount (LMR1S; Thorlabs, USA). Four planoconvex lens prescriptions of different focal lengths (*f*) were selected, based on commercial glass counterparts; *f* = +12.5 mm (37-385; Edmund Optics, USA), *f* = + 19.9 mm (LA1074; ThorLabs, USA), *f* = + 35.0 mm (37-791; Edmund Optics, USA), and *f* = + 49.8 mm (LA1213; ThorLabs, USA).

Following printing, the lenses were removed from the magnetic build plate and washed with neat isopropanol (10592921; FisherScientific, USA) for 9 minutes in a Mercury X washing station (Elegoo, China). The prints were removed and carefully air-dried using a compressed Ultra Pure Duster (Thorlabs, USA) before a post-print UV curing step for 20 minutes in a Mercury X curing station (Elegoo, China).

### Spin Coating and Lens Preparation

The printed lenses were rendered optically transparent by spin coating a thin layer of resin over both the curved and planar surfaces, minimising layer artefacts and surface structures from the printing process. For the convex surface, a spin coater (L2001A3-E463; Ossila, UK) was fitted with a custom 3D printed chuck with a 25 mm diameter well to accommodate a printed lens. The printed lens was cleaned again prior to coating with 100% isopropanol, airdried using compressed air, and placed in the chuck. 100 µl of Clear UV Resin (4^th^ Generation; VidaRosa, China) was deposited on the apex of the lenses (50 µl for *f* = + 49.8 mm to accommodate the shallower curvature) and spun for 10 seconds at 2000 rpm. The coated lenses were stored in darkened conditions for 30 minutes to allow the liquid resin to settle and were subsequently cured for 20 minutes using the Mercury X curer as described above.

For the planar surface, the spin coater was fitted with a 76 mm x 26 mm microscope slide chuck and a clean microscope slide placed in the chuck. A thin resin layer was created by depositing 100 µl of Clear Resin (v4; Formlabs, USA) on the slide and spinning for 10 seconds at 2000 rpm. The cleaned planar slide of the printed lens was carefully placed on to the resin-coated slide to avoid introducing air bubbles and was placed in a vacuum chamber (2 L,; Bacoeng, USA) fitted with a vacuum pump (Capex 8C; Charles Austen Pumps Ltd, UK). The assembly was maintained under a vacuum on 0.9 bars for 30 minutes before curing for 20 minutes as described above. The melded lens-slide combination was stored at -20°C for 3 minutes and carefully levered from the slide, relying on the differential thermal expansion of the glass and resin to remove the lens from the microscope slide.

### Spin-coat Thickness Measurements

To determine the thickness of the spin coated layers, coumarin-30, a non-polar green-emitting organic fluorophore, was prepared as a 10 mM stock in 100% isopropanol and mixed with 1 ml of Clear UV Resin (4^th^ Generation; VidaRosa, China) at a final concentration of 100 µM. Lenses from each of the four test prescriptions were spin coated and cured as described above.

The thickness of the fluorescent spin coated layer was measured by acquiring a 3D image stack using a confocal laser scanning microscope. An Olympus IX81 inverted microscope coupled to an FV1000 confocal laser scanning unit (Olympus, Japan) was used for imaging. Excitation of fluorescence was performed using a 488 nm argon laser (GLG3135; Showa Optronics, Japan) and fluorescence emission from coumarin-30 was detected by a photomultiplier tube (PMT) with a detection spectral window of 500 nm to 550 nm. Coated lenses were placed with the curved surface in contact with a Type 1.5 coverglass and imaged using a 10×/0.4 numerical aperture (NA) objective lens (Olympus, Japan). All images were acquired at the axial Nyquist sampling rate for the imaging objective (Δ*z* = 1.53 µm). The thickness of the fluorescent layer was measured using a linear plot profile of the fluorescence intensity in orthogonal (*x*,*z* and *y*,*z*) views of the 3D image stack using FIJI (v1.53t) [25]. Analysis was conducted using three replicate printed lenses with each focal length.

### White Light Interferometry and Stylus Profilometry

Methods for measuring the surface profile of planoconvex lenses include non-contact white light interferometry and contact stylus profilometry, with contact measurement approaches often only reporting the curvature of a linear trace instead of providing a 3D reconstruction of the specimen topology. Non-contact surface profiles were obtained using a white light interferometer (Wyko NT1100; Veeco Instruments Inc, USA) which used coherent light to generate interference fringes which are axially shifted through the optical surface, providing two-dimensional surface roughness and uniformity measurements. The interferometer was used in a vertical scanning interferometry configuration where an internal translator axially scanned in one direction during the measurement as the in-built camera detector recorded each frame. The non-contact approach provides an approximate 300 x 200 µm surface area measurement using a 20x objective, with a colour gradient to indicate height (Δ*z*) changes as well as read-out line profiles to provide sub-nanometre-scale surface roughness across the *x* and *y* axes of the specimen. The contact approach utilised a stylus profiler (Alpha-Step IQ; KLA Corp., USA) with 5 µm tip diameter for one-dimensional surface topology. The stylus profilometry technique provides millimetre-range one-dimensional measurements with sub-nanometre height resolution for curvature and roughness analysis.

### Interference Reflection Microscopy (IRM)

The underpinning theory of IRM relating to imaging of plano-convex lenses has been explained elsewhere [26,27]. Briefly, the printed lens specimens were placed convex side down on a Type 1.5 coverglass, bridging the stage insert of an IX81 inverted microscope coupled to an FV1000 confocal laser scanning unit (Olympus, Japan). An 80/20 beamsplitter was used in place of the dichroic filter in the confocal microscope, which facilitated a configuration to detect reflected light from the specimen plane. A 458 nm argon laser (GLG3135; Showa Optronics, Japan) provided incident light, which was reflected from refractive index boundaries at the specimen plane (i.e., coverglass-air and air-lens interfaces). Reflected light from each interface coincided, leading to constructive and destructive interference depending on the optical path difference of the two reflected beams (Figure 2a). The resulting image provided a 2D projection of the 3D topography of the specimen surface, where interference orders were separated along the optical axis. Equations 1 and 2 describe the axial separation of destructive and constructive interference orders, respectively [28,29], where *z* = fringe spacing, *N* = order, λ = wavelength of incident light, and *n_m_*= refractive index of the imaging medium.

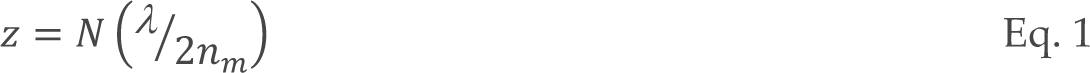

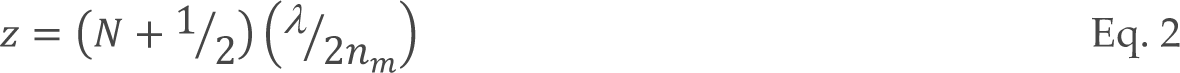

**Figure 2.**
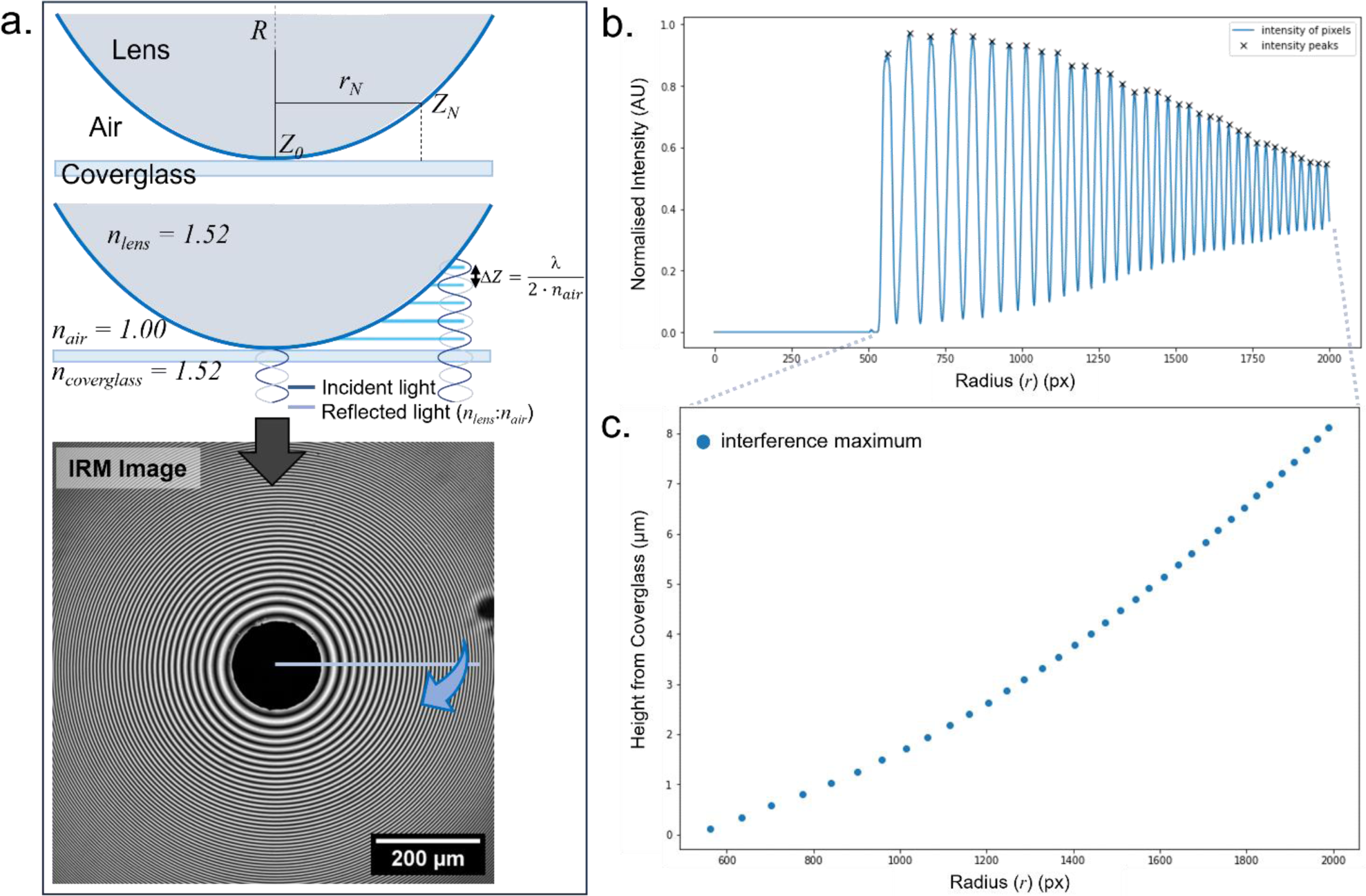
Reconstruction of Lens Curvature using Interference Reflection Microscopy (IRM). **(a)** A schematic of the principles of IRM (not to scale). Refractive index boundaries reflect incident light and constructive/destructive interference occurs depending on the relative distance between reflective boundaries **(b)** An IRM image of a lens was acquired, and a line intensity profile was measured. **(c)** The axial position of the interference maxima (Eq. 1) was plotted against the radial peak position from the line profile.

IRM images were acquired using a 10x/0.4 NA objective lens (Olympus, Japan) and the reflection signal was detected using a PMT with the detection limited to 458 nm ± 5 nm.

### Reconstructing 3D Surface Curvature and Quantifying Radius of Curvature from 2D IRM Image Data

All computational analyses of IRM data were performed using FIJI and Python 3.8.10 (64-bit) in a Spyder IDE 5.3.2 environment on an Elitebook 840 G7 (Hewlett-Packard, USA) running a 64-bit Windows 10 Enterprise operating system with an Intel^®^ Core™ i5-10310U 1.70GHz quad-core processor with 16 GB of 2666 MT/s DDR4 RAM.

The IRM image data were exported as .OIB files and pre-processed using FIIJ. The images were cropped to ensure the apex of the lens was centred in the image, and a median filter (σ = 2) was applied to remove any high-frequency noise in the data. Images were contrast adjusted using the Contrast Limited Adaptive Histogram Equalisation (CLAHE) plugin [30] (blocksize = 127, histogram bins = 256, maximum slope = 3.00) and converted to .PNG files for analysis.

A custom Python pipeline [31] was created to generate 3D reconstructions of the surface of the lens specimens from 2D IRM images and calculate the median radius of curvature for each lens. Briefly, the *Calibration* script was first used to verify the correct feature detection parameters for IRM data. The position of the zeroth order minimum was taken as the centre of the lens and the radius was noted as half the width of the image. The position of the intensity maxima along the radius was calculated using the *find_peaks* function, iteratively optimising the detection thresholds for peak height, distance, and prominence to ensure that each intensity maximum was detected. A line intensity profile noting the position of each interference maxima was generated along radius (*r*) (Figure 2b). The axial position of each interference maximum was calculated using Equation 1 and was plotted against distance to show the curvature of the lens (Figure 2c).

The surface curvature of each lens was reconstructed using the *3D Reconstruction* script with the optimised setting for each lens applied from the *Calibration* script. The 3D convex surface was reconstructed by assigning an axial position, as above, to each interference order detected along each radius (360 radii measured per image). The radius of curvature (*R*) value for each radius was calculated using Equation 3 [32].

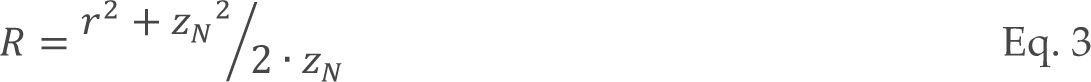

The measured radius of curvature for each lens was calculated using the *Radius Analysis* script in Python. The *R* value for each radius was compiled into a histogram that compared the experimental measurements to the theoretical *R* (i.e., the manufacturer-quoted value for the lens design file used for printed). The maximum peak position from each histogram determined the *R* value for each lens.

The measured and theoretical curvature data were plotted using Prism (v8.0.2) (GraphPad Software, USA).

### Tolansky Interferometry

Tolansky interferometry, a mode of multiple-beam interference imaging, was used as an alternative means to visualise the curved topology at the apex of the convex surface of the 3D printed lenses. The principle of Tolansky interferometry relies on two opposing highly reflective surfaces (in this case, an aluminium-coated coverglass and aluminium-coated 3D printed lens) which generate multiple reflected beams that undergo constructive and destructive interference [33–35]. The multiple beam combination acts to modify the Haidinger rings formed at the focus of the objective lens such that the nodal spacing is not altered, but the intensity distribution of the interference orders is changed. This effectively increases the axial resolution compared with IRM, such that topological features as small as 5 Å, or better, can be resolved within the interference maxima [28]. Moreover, uncoupling the reflective specimen from the reflective coverglass provides a means to translate the interference orders through the optical axis, in turn ‘scanning’ the topography of the convex lens surface in a way that IRM cannot.

A 3D printed lens (*f* = + 19.9 mm) and a Type 1.5 coverglass were vapour coated with a thin layer of aluminium using a thermal evaporator coating system (E306A; Edwards Vacuum, UK). Briefly, a small quantity of aluminium foil was heated on a tungsten filament under vacuum (1.0×10^-5^ Pa), depositing vapourised aluminium on the surface of the lens and coverglass.

A custom steel objective lens collar was fabricated and fitted to a 10×/0.4 NA objective lens (Olympus, Japan). The aluminium-coated coverglass was bonded to the top of the collar using a thin layer of epoxy resin around the circumference of the objective lens housing. An adjustment screw was included in the collar to facilitate the positioning of the collar-coverglass assembly relative to the focal length of the objective lens (approximately 3.1 mm). The aluminium-coated lens specimen was suspended over the stage insert of an inverted IX81 microscope coupled to a confocal laser scanning unit (Olympus, Japan). Two glass microscope slide spacers were inserted to raise the test lens so that the modified objective underneath could be focussed near to the lens surface. Tolansky interferometry was performed using the same reflection setup as in IRM experiments but employed z-scanning which uncoupled the specimen from the coverslip. Altering the relative distance between the coverglass and lens specimen resulted in axial translation of interference orders and provided a means of visualising the curved apical surface by merging images acquired at different axial positions into a single hyperstack colour-coded by depth using FIJI.

### Optical Transmission Measurements

The percentage transmission of the printing resin was measured by comparing the mean intensity of transmitted light through resin blocks of varying thickness compared to the transmission through a single Type 1.5 coverglass. Resin blocks of thickness from 1 mm to 6 mm were printed using the optimised printing parameters used for lens printing. The blocks were placed on a Type 1.5 coverglass and imaged using an IX81 inverted microscope coupled to an FV1000 confocal laser scanning unit (Olympus, Japan) configured to detect scanned transmitted light. The transmissivity of unprocessed and processed blocks (i.e., naïve and spin coated, respectively) was measured using three discrete wavelengths across the visible spectrum sequentially; a 458 nm argon laser, a 515 nm argon laser, and a 633 nm helium-neon laser (GLG3135; Showa Optronics, Japan). Images were acquired using a 4×/0.1 NA objective lens (Olympus, Japan), with dimensions 64 × 64 pixels and Kalman averaging (n = 5 frames) to minimise contributions to the image from print structures. The mean intensity across the field was measured using FIJI and compared to the optical throughput of the coverglass alone. Linear fits were conducted using Prism.

### Beam Profilometry

An optical setup was constructed to measure the focusing ability of 3D printed lenses compared to their commercial glass counterpart. A complete parts list is included in the Supplementary Material. A 633 nm helium-neon laser source with an initial beam diameter measuring 600 µm was passed through a neutral density filter and was steered using two gimbal-mounted mirrors. The beam was expanded to a final diameter of 12.5 mm using two sequential beam expanders, first through a 2.5× beam expander and then through a 7.5× beam expander. The 3D printed lens was mounted in a fixed mount (LMR1S/M; Thorlabs, USA) and a dual scanning slit beam profiler was mounted on a linear translational stage to facilitate movement along the optical axis to map the beam diameter with respect to post-lens propagation distance. Perpendicular measurements (*x* and *y* axes) of the focused beam diameter (^1^⁄_𝑒2_) were measured at increments along the optical axis and compiled to provide a beam profile for three replicates of various planoconvex lens prescriptions. The ^1^⁄_𝑒2_ beam waist radius (*w_0_*) was calculated from the measured beam diameter and the Rayleigh Range (*z_R_*) for each lens was calculated using Equation 4 [36].

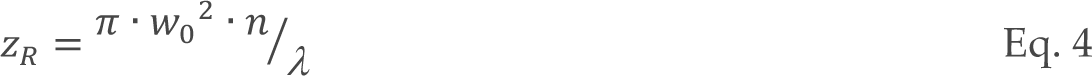

### Brightfield Transmission Microscopy

A brightfield transmission microscopy setup was constructed to demonstrate the performance of a 3D printed lens in an imaging setup. A complete parts list is included in the Supplementary Material. A 10 mm stage micrometer with 50 µm intervals (R1L3S1P; ThorLabs, USA) and a thin section of linden tree stem (*Tilia europaea*) were imaged to assess field of view and to determine the resolution of the system. A blue light-emitting diode (LED) source (λ = 470 nm) (M470L2-C1; ThorLabs, USA) was used to illuminate the specimen. Light from the LED was brought to the specimen using a 3D printed *f* = + 49.8 mm planoconvex condenser lens (modelled on LA1213; ThorLabs, USA) and transmitted light was detected using a monochrome complementary metal-oxide-semiconductor (CMOS) camera (DCC3260M; ThorLabs, USA). Image acquisition was controlled via ThorCam (64-bit, v3.7.0) (ThorLabs, USA).

## Results

### Surface Characterisation of 3D Printed Lenses

#### Measuring the Thickness of Spin Coated Resin Layers on 3D Printed Lenses

The thickness of the spin-coated resin layer on the printed convex lens surface was measured using 3D confocal laser scanning microscopy. The coumarin-30-spiked VidaRosa Clear resin was spin coated onto the convex surface using the same spin settings as for other lenses, creating a fluorescent resin layer (Figure 3a). A 3D confocal z-stack visualised the spin-coated layer relative to the lens surface (not fluorescent, ergo dark) (Figure 3b). The mean spin coated thickness of the fluorescent resin was measured for three replicates of various lens prescriptions, with the spin coat thickness routinely ranging from 25 µm to 45 µm (mean thicknesses; *f*12.5 mm = 30.00 µm, *f*19.9 mm = 41.31 µm, *f*35.0 mm = 42.00 µm, *f*49.8 mm = 28.50 µm) (Figure 3c).

**Figure 3.**
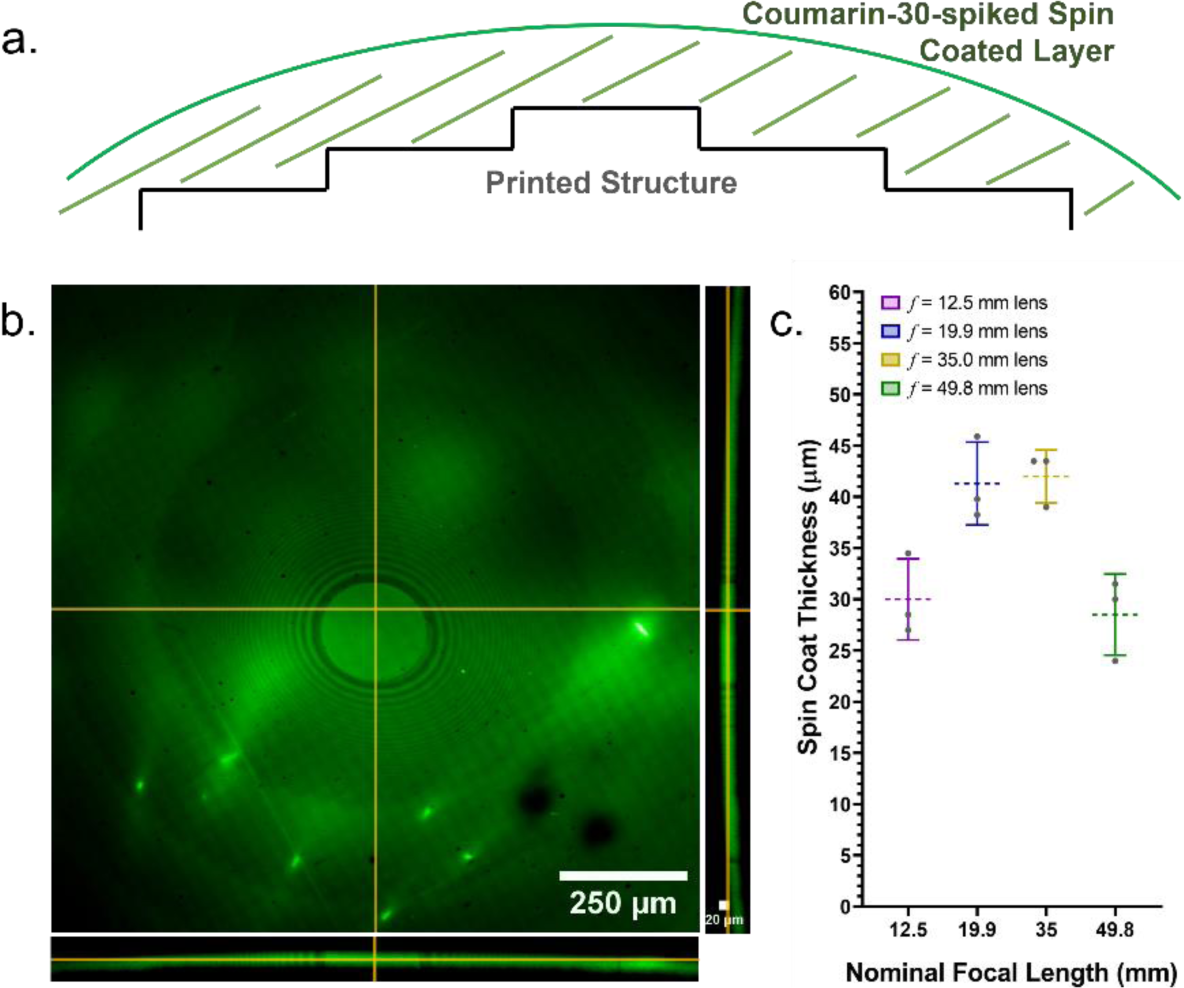
Thickness Measurements of the Spin-Coated Surface Layer of 3D Printed Lenses. **(a)** A schematic of the spin-coated layer spiked with the organic dye, Coumarin-30 (not to scale) **(b)** An average intensity projection and orthogonal views of the spin-coated layer of a 3D printed lens. **(c)** Comparison of the spin coated layer thickness across three replicate 3D printed lenses for four lens prescriptions (*f* = + 12.5 mm, purple; *f* = + 19.9 mm, blue; *f* = + 35.0 mm, yellow; *f* = + 49.8 mm, green). The lenses had a median coat thickness ranging from 28.5 µm to 42.0 µm.

#### Comparing the Surface Curvature and Uniformity between 3D Printed and Commercial Glass Lenses

The surface curvature of 3D printed lenses was first measured by conventional means using a commercial white light interferometer and a stylus profilometer. However, white light interferometry usually requires higher reflective surfaces for accurate surface measurements, and stylus profilometry is typically restricted by measuring only orthogonal straight lines along the *x* and *y* axes of the lens (Supplementary Figure 2). An alternative method was required that accurately reconstructed the transparent three-dimensional surface of the printed lenses, which provided a robust method to identify surface curvature defects that could impact optical performance.

Using the methods outlined in Figure 2, 2D IRM image data of printed planoconvex lens surfaces (Figure 4a) were processed to create 3D renders of the curved surface (Figure 4b). The radius of curvature was measured for each radius around the circumference of the lens and plotted as a histogram to calculate the median radius of curvature for each printed lens (Figure 4c). The radii of curvature for three replicate printed lenses of four prescriptions were compared to commercial glass lenses (Figure 4d, Supplementary Figure 3). The radius of curvature of 3D printed lenses concurred with their glass counterparts, with a slight increase due to the additive spin coating process. However, this did not hold true for longer focal length lenses, where the increased radius of curvature is more pronounced for longer focal length lenses with larger variation due to the shallower curvature. The mean radius of curvature (± standard deviation) measured; *R* (*f _+_* 12.5 mm) = 10.76 mm ± 1.04 mm, *R* (*f _+_* 19.9 mm) = 11.31 mm ± 0.66 mm, *R* (*f _+_* 35.0 mm) = 18.83 mm ± 0.40 mm, *R* (*f _+_* 49.8 mm) = 31.44 mm ± 4.01 mm. These values compare to the theoretical radii of curvature for their glass counterparts; *R* (*f _+_* 12.5 mm) = 9.80 mm, *R* (*f _+_* 19.9 mm) = 10.30 mm, *R* (*f _+_* 35.0 mm) = 18.10 mm, *R* (*f _+_* 49.8 mm) = 25.80 mm.

**Figure 4.**
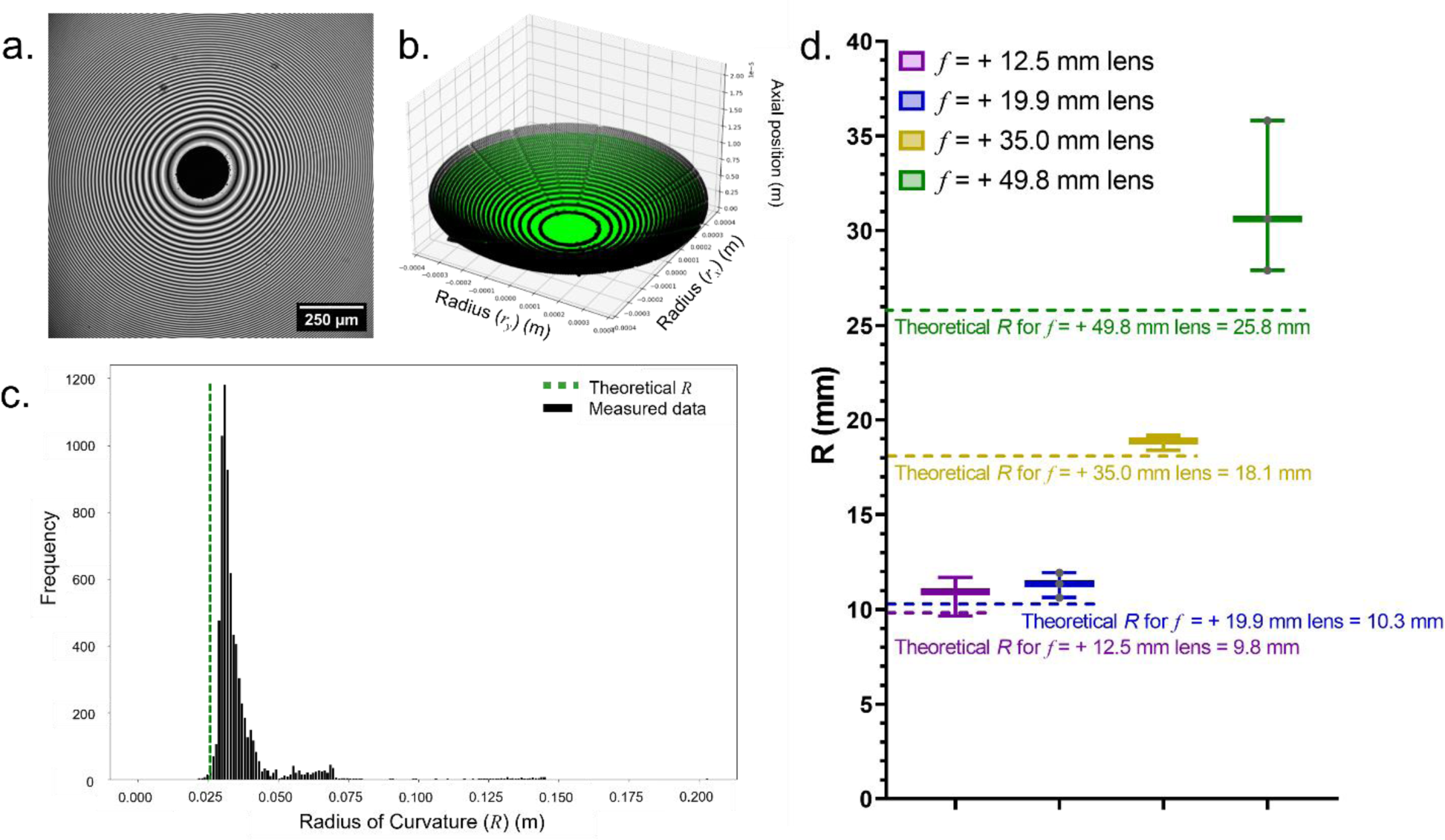
Radius of Curvature Measurements of 3D Printed Lenses using IRM. **(a-b)** Surface reconstruction of a 3D printed lens from 2D IRM data using method presented in Figure 2. **(a)** An IRM image of a 3D printed lens surface (λ = 458 nm). **(b)** A 3D reconstruction of the printed lens surface (black datapoints) compared to the theoretical curvature from the print design (green). **(c)** A histogram of the distribution of radius of curvature (*R*) values measured around the circumference of the lens (black) compared to the theoretical *R* (green) (See Eq. 2). **(d)** Measured radius of curvature values of three replicate 3D printed lenses for lens prescriptions (*f* = + 12.5 mm, purple; *f* = + 19.9 mm, blue; *f* = + 35.0 mm, yellow; *f* = + 49.8 mm, green).

The IRM images of 3D printed lenses routinely featured a larger than expected zeroth order minimum. This suggested that the apex of the convex lenses was flat, but this was not observed in white light interferometry or stylus profilometry experiments (Supplementary Figure 2). Moreover, the curvature of the zeroth order area measured using these methods agreed with the theoretical curvature and the median radius of curvature measured by IRM. To conclude that the zeroth order area was curved, a modified *z*-scanning Tolansky interferometry method was employed (Figure 5a). Uncoupling of the lens specimen from the objective setup permitted translation of the interference orders over the lens surface in unison with the axial movement of the modified objective housing (Figure 5b). A maximum intensity projection from a Tolansky interferometry z-series revealed the nanoscale surface profile and the continuous curved surface of the printed lens (Figure 5c), confirming the observations using white light and stylus profilometry.

**Figure 5.**
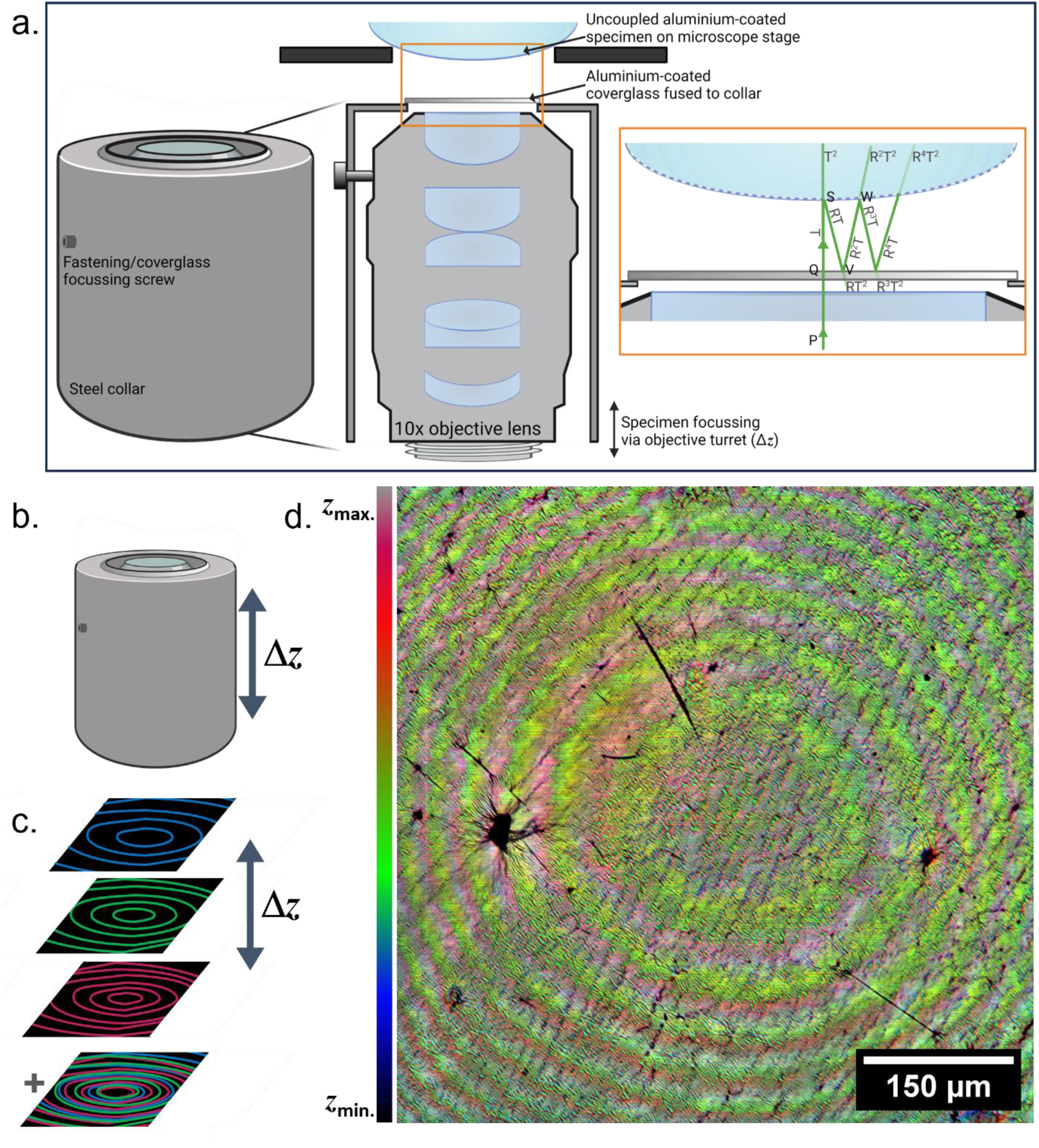
Tolansky Interferometry Confirms the Apical Surface Curvature of 3D Printed Lenses. **(a)** A schematic (not to scale) describing the principles of Tolansky interferometry and the design of a custom objective lens to permit 3D measurements of surface curvature of an aluminium-coated specimen. **(b)** Axial translation of the objective mount results in **(c)** translation of the interference orders, which can be colour-coded by depth and merged into a *z*-projection of a Tolansky interferometry acquisition. **(d)** The concentric interference maxima from each axial position are false coloured according to their depth and super-imposed, revealing the curved surface and nanoscale topology of the apical lens surface with higher resolution than interference optical microscopy can provide.

### Optical Characterisation of 3D Printed Lenses

#### Comparing the Transmissivity of 3D Printed Resin to Commercial Glass Lenses

The transmissivity of the 3D print resin substrate was measured to verify the optical properties of the clear resin. The mean intensity of the transmitted light was normalised compared to a control of the same intensity of light passing through a Type 1.5 coverglass (Figure 6). Block transmissivity was increased by approximately 2.25× up to greater than 90% across all tested wavelengths following spin-coating, comparable to uncoated N-BK7 glass often used to manufacture glass bulk optics [37].

**Figure 6.**
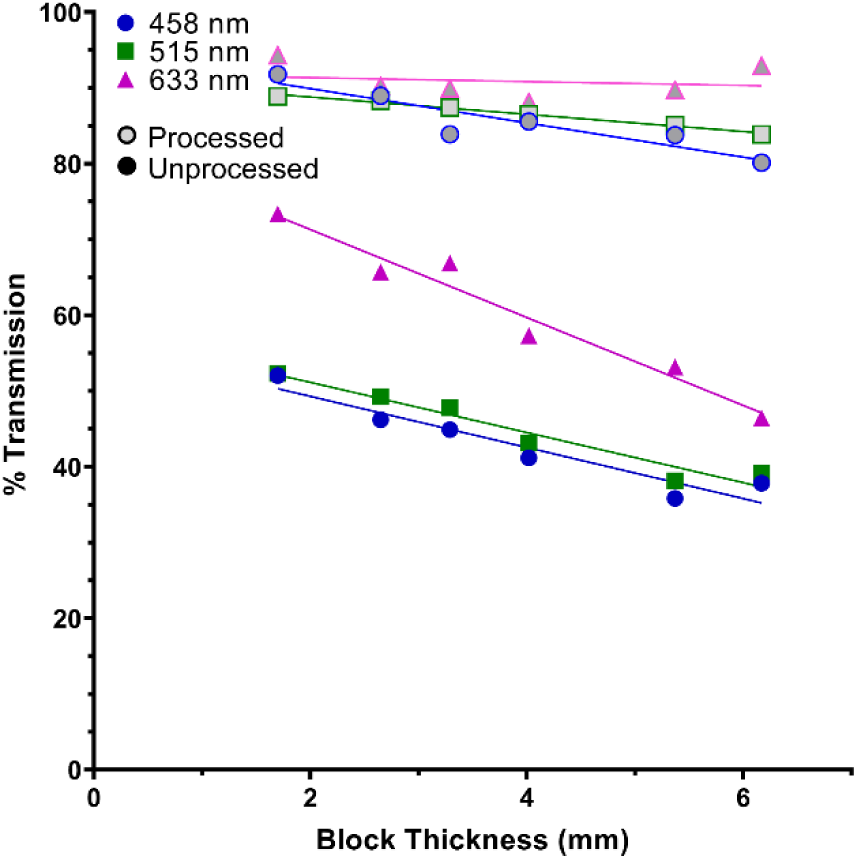
Optical Throughput of Naïve and Post-processed Resin Blocks. Naïve (i.e., unprocessed) blocks (solid markers) exhibited lower optical throughput than processed blocks (hollow markers). Optical throughput was typically higher with longer wavelengths of transmitted light, with processed blocks achieving up to 94.33% transmission at a wavelength of 633 nm.

#### Measuring the Optical Performance of 3D Printed Lenses

The optical performance of four 3D printed planoconvex lens prescriptions was measured in triplicate using the setup shown in Figure 7a. The focal length approximately matched the theoretical values for all lenses, except for *f* = 49.8 mm (Figure 7). The measured focal lengths for each lens prescription were *f*12.5 mm = 13.5 mm, *f*19.9 mm = 19.0 mm, *f*35.0 mm = 35.0 mm. These data presented an error of *f*12.5 mm = 8.0%, *f*19.9 mm = -4.5%, and *f*35.0 mm = 0.0% compared to the focal lengths of their glass counterparts. The longer theoretical focal length lenses (i.e., *f*49.8 mm) did not focus the light as expected. The beam profile was elongated along the optical axis at the beam waist and was significantly displaced from the theoretical focal length, indicating the presence of spherical aberration. Each of the printed lenses were compared to the focusing performance of their commercial glass counterparts using the same profiling setup. The glass commercial lenses performed as expected, resulting in the correct focal length for each prescription.

**Figure 7.**
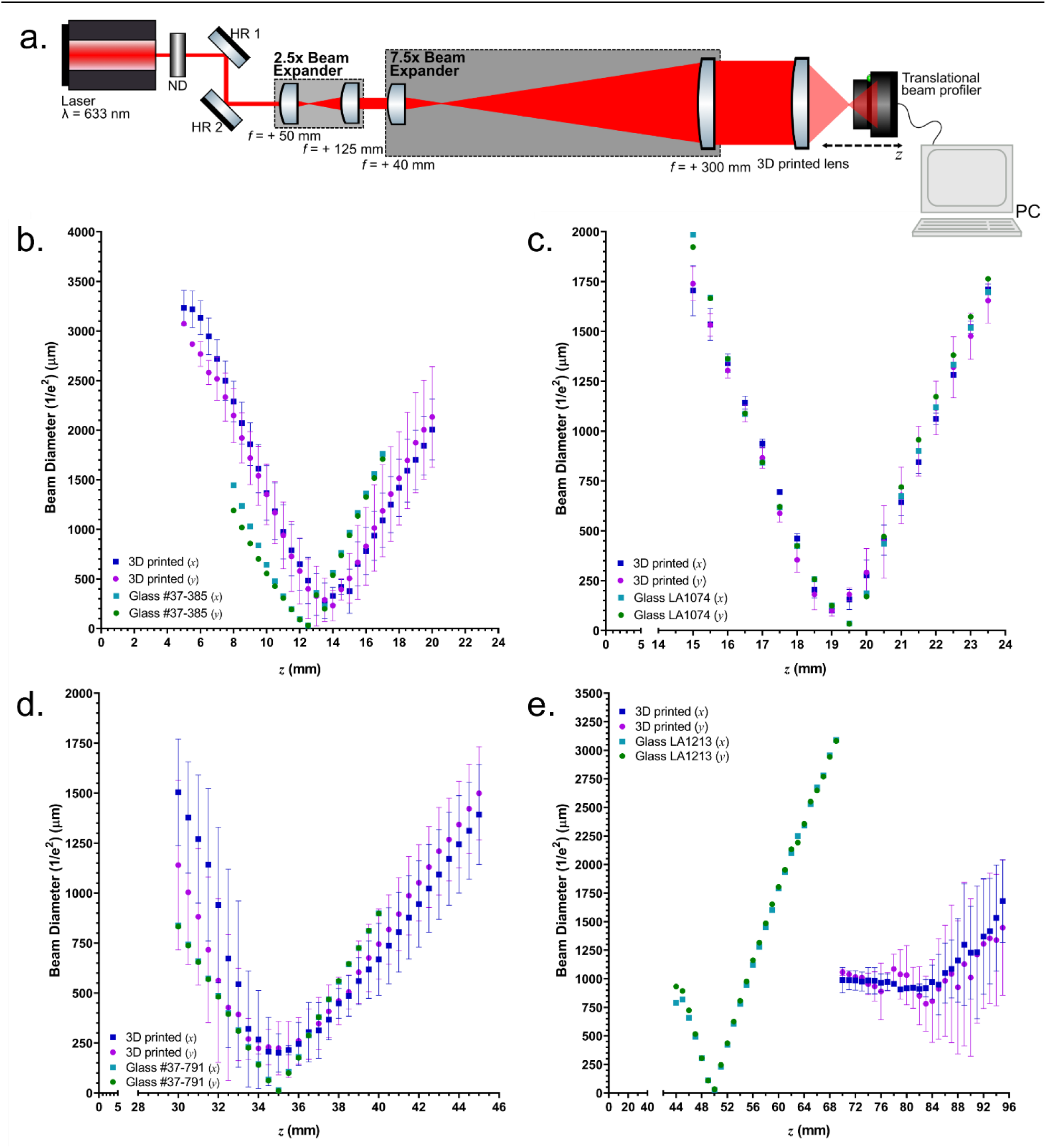
Comparing the Focusing Performance of 3D Printed Lenses. (a) An optical setup to measure the beam profile of a 633 nm laser focused by a 3D printed lens. Two concurrent beam expanders provide a total beam expansion of 18.75×, creating a 12.5 mm diameter beam that uses the full numerical aperture of the printed lenses. A beam profiler is translated along the optical axis to measure the focal length and metrics of the focused beam. ND = Neutral Density Filter, HR = High Reflector (Mirror), all lenses shown are commercial glass planoconvex lenses, save for the final 3D printed lens under observation. **(b-e)** The beam diameter along the optical axis of several 3D-printed lens prescriptions is presented. Both the *x* (blue) and *y* (purple) beam diameters are noted as a function of 1/e^2^. The beam profiles for lenses of theoretical focal length **(a)** + 12.5 mm, **(b)** + 19.9 mm, **(c)** + 35.0 mm, and **(d)** + 49.8 mm are presented. The focal lengths of (a), (b), and (c) concurred with their theoretical glass counterparts (presented in green for illustration in (b) and (d)) but did not agree for longer focal length lenses in (d).

The beam waist (*w_0_*) and Rayleigh Range were measured for each 3D printed lens and were compared to the commercial glass counterpart, except for the *f*49.8 mm lenses due to the severity of the optical aberrations (Table 1). The mean *w_0_* and the Rayleigh Range values for the lens prescriptions are noted in Table 1. Although the focal length of these 3D printed lenses conformed with their commercial glass counterparts, the mean *w_0_* value was routinely larger with 3D printed lenses. Moreover, the Rayleigh range was also increased proportionally to the enlarged *w_0_*. However, 3D printed lenses were ultimately able to focus a beam to a discrete point which demonstrated promise for implementation in optical systems.

**Table 1.** Comparison of Beam Parameters in 3D printed and Glass Lenses. The mean beam waist radius (*w_0_*) was measured for three different printed and glass planoconvex lens prescriptions, along with the standard deviation (SD) for three replicate printed lenses. The Rayleigh range (*z_R_*) was calculated for each prescription based on the mean *w_0_* value.

| Lens Prescription | $f_{12.5 \text{ mm}}$ | | $f_{19.9 \text{ mm}}$ | | $f_{35 \text{ mm}}$ | |
| --- | --- | --- | --- | --- | --- | --- |
|  | 3D Printed | Commercial Glass | 3D Printed | Commercial Glass | 3D Printed | Commercial Glass |
| Mean beam waist radius ($w_0$) ($\mu\text{m}$) (± SD) | 62.40 ± 21.55 | 16.25 | 25.26 ± 5.06 | 17.46 | 56.67 ± 24.10 | 6.39 |
| Rayleigh range ($z_R$) (mm) | 19.32 | 1.31 | 3.17 | 1.51 | 15.93 | 0.20 |

#### Using 3D Printed Optics for Brightfield Transmission Imaging

The imaging performance of 3D printed optics was tested using the setup presented in Supplementary Figure 5, where a 3D printed lens (*f* = + 49.8 mm) was used as the condenser in a brightfield transmission microscope. The setup resulted in a field of view measuring approximately 2.3 mm wide with high contrast across the full field (Figure 8a). Moreover, brightfield transmission imaging was demonstrated using a cross section of a linden tree stem, resolving the intricate differentiated tissue layers and structures on the order of 6 µm (Figure 8b). This sub-cellular resolution demonstrates the potential of 3D printed optics in biological imaging systems.

**Figure 8.**
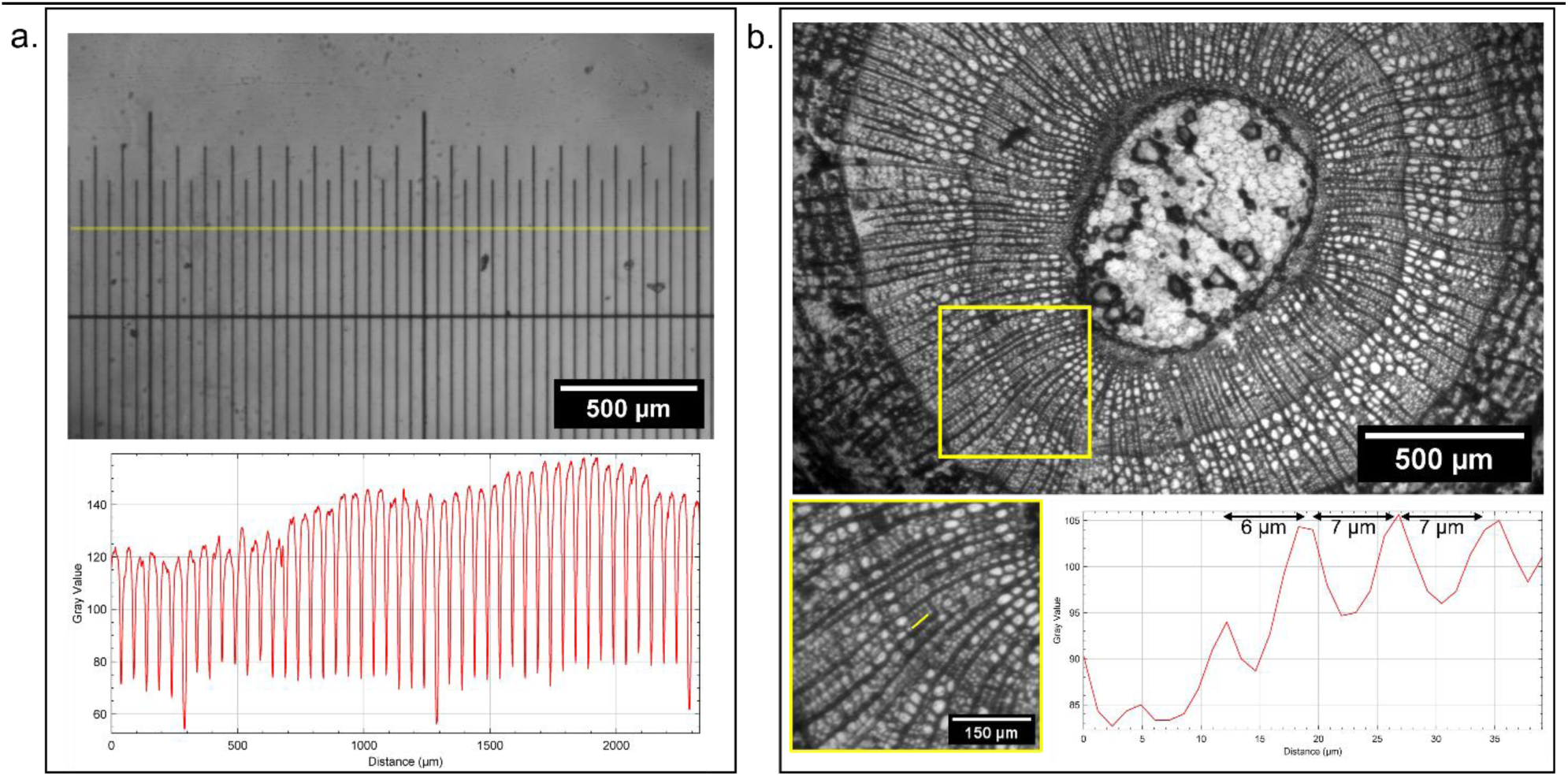
Brightfield Transmission Imaging Using a 3D Printed Condenser Lens. Images demonstrating the application of a 3D printed condenser lens (*f* = + 49.8 mm) **(a)** An image of a stage micrometer (R1L3S1P, Thorlabs, USA) measuring a field of view of approximately 2.3 mm. A line intensity profile (averaged over 5 pixels thickness) is plotted against distance, showing high contrast across the field of view. **(b)** An image of a linden tree (*Tilia europaea*) stem showing the intricate tissue layers. A magnified region of interest is presented with a yellow box, and this is digitally magnified to show individual plant cells. A line intensity profile (averaged over 5 pixels thickness) is plotted against distance and demonstrates resolution at a level that would be sufficient to record subcellular detail.

## Discussion

We have demonstrated robust, repeatable, and accessible methods to manufacture planoconvex lenses using 3D printing with consumer-grade instrumentation and printing materials. Moreover, we have characterised and compared the quality of 3D printed lenses against commercially available glass counterparts of the same prescription. A spin-coating method was employed to obviate the stepped print structure and to render the printed lenses smooth and transparent, resulting in a thin surface coating on the order of 35 µm thick. White light and stylus profilometry were used to assess the surface quality of printed lenses, while confocal IRM was used to reconstruct the surface topology of 3D printed lenses with high axial resolution and quantify the radius of curvature, which matched that of glass lens counterparts. It is important to note that, while IRM in confocal scanning mode provides increased contrast due to coherent illumination and the rejection of out-of-focus light by a pinhole aperture, widefield IRM also presents feasible and accessible method to achieve similar, albeit lower-contrast, IRM data. Tolansky multiple beam interferometry corroborated the profilometry data by revealing the curved topology of the apex of the convex lenses. The printing and processing steps facilitated comparable optical throughput to glass lenses, with greater than 90% transmission across the visible spectrum.

The optical performance of 3D printed lenses was determined by measuring their ability to focus a beam of light. Shorter focal length printed lenses were observed to have the same focal length as their glass counterparts, with moderately increased beam diameters and Rayleigh ranges. The performance of 3D printed lenses was limited in longer focal length printed lenses (i.e., *f* = + 49.8 mm), which exhibited extended focal lengths indicative of spherical aberration. However, our findings present 3D printing as a viable option for the manufacturing of high-quality lenses for optical instrumentation and rapid prototyping as a less costly and more accessible alternative to bulk glass optics. Finally, we demonstrated the use of 3D printed lenses for brightfield transmission imaging. Despite the aberrations observed during beam profilometry experiments using printed *f* = + 49.8 mm lenses, the imaging results were promising. Sub-cellular spatial resolution was achieved with high contrast over a 2.3 mm field of view, showing great promise for the use of 3D printed optics in microscopy.

Previous studies developing 3D printed optics have used a variety of additive manufacturing methods, these with most of these typically being costly, requiring specialist equipment, or being mainly focussed on using consumer-grade printers to produce micro-optics. Fused deposition modelling, where thin glass filaments are melted, extruded and cooled into the shape of a lens, has resulted in 3D printed glass bulk optics [38]. However, the silicon dioxide substrate requires careful mixing with titanium dioxide and a complex series of drying, burnout and sintering steps performed at over 1000 °C that limit users without access to specialist equipment. Alternative filament-based methods have used CO_2_ lasers to print transparent glass lenses from a single mode optical fibre and fused quartz filaments [39–41]. However, these molten glass methods often result in layering defects that reduce optical performance.

Stereolithography approaches have resulted in various techniques to manufacture resin-based optics. These methods routinely use a two-photon polymerisation-based platform to manufacture microlens arrays (MLAs) and optics, although print sizes are typically limited to only a few mm in diameter. Moreover, two-photon-based instrumentation can be prohibitively costly and presents a barrier to entry for accessible 3D printed optics. Printed lenses using, for example, a Nanoscribe printer have recently been produced [42] but they retain microstructures and layering artefacts resulting from the printing process and curvature defects that ultimately impact on their optical performance. Moreover, these techniques are often limited by their small print sizes and high materials and instrument costs, somewhat restricting their use to MLAs and other micro-optics. Recent developments in two-photon microprinting have successfully been used to fabricate bespoke micro-optics, such as 30 µm-diameter Fresnel elements for x-ray microscopy, using photopolymerising resins on a supportive silicon nitride membrane [43]. Foveated compound microlenses have also recently been produced using femtosecond direct laser writing [18], but all two-photon methods require a costly and complex ultrashort pulsed laser source. Microlens arrays have been printed using UV-induced photopolymerisation, with recent improvements seeing expansion of MLAs over large flexible substrates to improve optical performance [44] and the introduction of vibrating projection lenses during printing to smooth the surface of 3D printed micro-optics [45]. Aspheric lenses have been manufactured using UV photopolymerisation, however these specialised lenses required assembly with corrective quartz substrates and refractive index matching liquids in order to focus light, but suffered from both chromatic and spherical aberration [46]. Each of these 3D printing methods are based on inaccessible and specialist equipment, which often produce 3D printed lenses that do not compare to the performance of their commercially produced glass counterparts.

We have described an accessible, low-cost, and reproducible method for manufacturing bespoke 3D printed lenses and a suite of characterisation methods that demonstrate their comparable performance to glass lenses. We implemented profilometry methods centred on IRM that provide high-resolution topographical reconstructions of the lens geometry, which offer alternative analysis methods for IRM and standing wave microscopy [26,47]. Using 458 nm illumination for IRM and 633 nm light for beam profiling ensured that the material properties of the printed lenses did not change during observation, as the resin absorbance peak is noted as 405 nm. The Tolansky interferometry mapping, together with white light and stylus profilometry data, suggest that the enlarged zeroth order present in IRM images was an inherent interference artefact, perhaps contributed by subtle refractive index differences between the printed layers caused by compounding exposure to light during printing [48,49]. However, despite this apparent interference artefact across all lenses, it did not impact on their optical performance. Short focal length lenses with inherently higher curvatures performed in line with their commercial glass counterparts, but longer focal length lenses on the order of *f* = 50 mm were subject to optical aberrations. This suggests that future applications creating 3D printed multi-lens systems would better suit the inclusion of shorter focal length elements. Overall, the beam waist radius and Rayleigh range were increased compared to glass lenses, however this did not impact the ability of 3D printed lenses to effectively focus light.

The potential for additive manufacturing for bespoke optics, rapid design prototyping, and field diagnostics is huge. Our data show that high-quality optical elements can be produced at low cost with consumer grade equipment, totalling approximately £300. The only additional outlet would be a spin coater, which can be procured for less than £1,000. The total cost in producing a single 3D printed lens was approximately £0.11, as opposed to upwards of £50 for commercial high-grade lenses. The demonstrated optical performance of 3D printed lenses shows great promise for optical imaging and prototyping optical instrumentation. Moreover, we have shown separately that 3D printed lenses can be implemented in bioimaging applications, using both absorption and fluorescence imaging modalities [50]. The potential impact of these accessible and open manufacturing methods could also impact across low resource settings for rapid diagnostics of blood smears, for example, where 3D printing has already made significant impacts. The combination of previous 3D printed microscope chassis with 3D printed lenses would be a natural evolution to produce the first fully 3D printed optical microscope.

## Conclusions

We present an accessible 3D printing method to manufacture high-quality optical lenses and provide characterisation methods to quantify their performance. With the prohibitive cost of bespoke bulk glass optics and difficulties in their manufacture, 3D printing offers a viable method to produce a range of lenses with a high degree of reproducibility. The quality of 3D printed lenses was determined by comparing their surface curvature, optical throughput, and ability to focus light compared to their commercial glass counterparts. Glass and 3D printed lenses were observed to behave similarly for a range of short focal length prescriptions, but longer focal lengths introduced a high degree of spherical aberration. The same trend was true for the surface curvature, where highly curved lenses conformed to the radius of curvature of their glass counterparts, while the surface coat thickness was routinely on the order of 35 µm. The transmissivity of 3D printed lenses was comparable to that of bulk N-BK7 glass across the visible spectrum. Moreover, 3D printed lenses were implemented in a brightfield transmission microscopy setup and facilitated high-quality imaging that demonstrated promise for future applications. Each of these observations concluded that 3D printing is a viable approach to reproducibly producing large volumes of high-quality optical elements that provide promise for prototyping, imaging applications, and field diagnostics.

## Additional Information

### Acknowledgements

The Authors would like to thank Dr Ross Scrimgeour (Institute for Cancer Research, UK) for his helpful comments on the IRM analysis code, Mr Rian MacDonnchadha (University of Glasgow, UK) for his insights on 3D printing instrumentation, and Miss Kay Polland for her assistance with figure preparation.

The schematics and workflow presented in Figure 1 and Figure 5 were prepared using <u>BioRender.com</u> (Licence Number: SR25JKZ3PC).

LMR, SF, and WBA were funded by the Leverhulme Trust. JC and RB were funded by the Engineering and Physical Sciences Research Council (grant EP/S032606/1, studentship EP/T517938/1) and the UK Royal Academy of Engineering (Engineering for Development Fellowship scheme RF1516/15/8). BW was funded by a Royal Microscopical Society Summer Studentship. LC and LDW were funded by an EPSRC iCASE studentship (EP/Y528833/1). GM was funded by The Medical Research Council, MR/K015583/1, and the Biotechnology and Biological Sciences Research Council, BB/P02565X/1 and BB/T011602/1.

### Data and Code Availability

The analysis code used to reconstruct and quantify IRM data is openly available on GitHub (<u>github.com/Liam-M-Rooney/IRM_lenses)</u>. All data underpinning this publication are openly available from the University of Strathclyde KnowledgeBase at <u>doi.org/10.15129/bc849aae-198c-45f3-9e7f-2613532726af</u>.

For the purpose of open access, the authors have applied a Creative Commons Attribution (CC BY) licence to any Author Accepted Manuscript version arising from this submission

### Authors’ Contributions

LMR: conceptualisation, data curation, analysis, investigation, methodology, supervision, validation, visualisation, manuscript preparation and review.

JC: conceptualisation, analysis, investigation, methodology, validation, visualisation, manuscript preparation and review.

BW: analysis, investigation, methodology, validation, software, visualisation, manuscript review.

YSK: analysis, investigation, methodology, validation, visualisation, manuscript review.

LC: investigation, methodology, manuscript review.

LDW: investigation, methodology, manuscript review.

SF: methodology, validation, manuscript review.

WBA: conceptualisation, methodology, manuscript review.

RB: conceptualisation, funding acquisition, project administration, supervision, manuscript review.

GM: conceptualisation, funding acquisition, project administration, supervision, manuscript review.

