## Supplementary Material for "Printing, Characterising, and Assessing Transparent 3D Printed Lenses for Optical Imaging"

---

#### Supplementary Information

##### Optimised 3D Printing Parameters

3D printing was conducted using the following parameters.

**Burn-In Layers**

- Number of Layers – 10
- Exposure Time – 45 s
- Transition Layers Count – 0
- Light-off Delay – 0 s
- Lift Distance – 0 mm
- Lift Speed – 65 mm/m
- Retract Speed – 150 mm/m
- Light Intensity – 100%

**Normal Layers**

- Layer Thickness – 10 μm
- Exposure Time – 10 s
- Lift Distance – 0 mm
- Lift Speed – 65 mm/m
- Retract Speed – 150 mm/m
- Light-off Delay – 10 s
- Light Intensity – 100%

**Scene Scale Compensation**

- Scale X – 100%
- Scale Y – 100%
- Scale Z – 100%

#### Code Availability

The analysis scripts to reconstruct the 3D surface of printed lenses using IRM data and calculate their radius of curvature are available at github.com/Liam-M-Rooney/IRM_lenses.

#### Beam Profiler Setup Parts List

The parts required for the setup detailed in Figure 7a are listed below. All parts were procured from Thorlabs, USA.

- HeNe Laser, 632.8 nm, 10 mW, Polarized, 100-240 VAC - HNL100LB
- Mounted Continuously Variable ND Filter – NDC-100C-2M
- 2x Broadband Dielectric Mirror, 400 - 750 nm – BB1-E02
- 2x Kinematic Mirror Mount for Ø1" Optics – KM100
- Ø1" (Ø25.4 mm) N-BK7 Plano-Convex Lens, $f$ = 49.8 mm – LA1131
- Ø1" (Ø25.4 mm) N-BK7 Plano-Convex Lens, $f$ = 124.6 mm – LA1986
- Ø1" (Ø25.4 mm) N-BK7 Plano-Convex Lens, $f$ = 39.9 mm – LA1422
- Ø1" (Ø25.4 mm) N-BK7 Plano-Convex Lens, $f$ = 299.0 mm – LA1484
- 5x Ø1" Lens Mount with Internal and External SM1 Threads, M4 Tap – LMR1S/M
- Dual Scanning Slit Beam Profiler, 200 - 1100 nm, Ø2.5 µm - Ø9 mm, Metric - BP209-VIS/M

#### Brightfield Transmission Microscope Setup Parts List

The parts required for the setup detailed in Supplementary Figure 5 are listed below. All parts were procured from Thorlabs, USA.

- Blue (470 nm) Collimated LED for Olympus BX & IX, 1600 mA – M470L2-C1
- T-Cube LED Driver, 700 mA Max Drive Current – LEDD1
- LED Connection Cable, 2 m, M8 Connector, 4 Wires – CAB-LEDD1
- Zero Aperture Iris, Ø50.0 mm Max Aperture – D50SZ
- 2x Ø1" (Ø25.4 mm) N-BK7 Plano-Convex Lens, $f$ = 74.8 mm – LA1608
- 2x Ø1" (Ø25.4 mm) N-BK7 Plano-Convex Lens, $f$ = 34.9 mm – LA1072
- XY Mount for 1/2" - 3" Rectangular Optics, M4 Taps – XYF1/M
- High-Resolution USB 3.0 CMOS Camera, 1936 x 1216, Global Shutter, Monochrome Sensor – DCC3260M

### Supplementary Figures

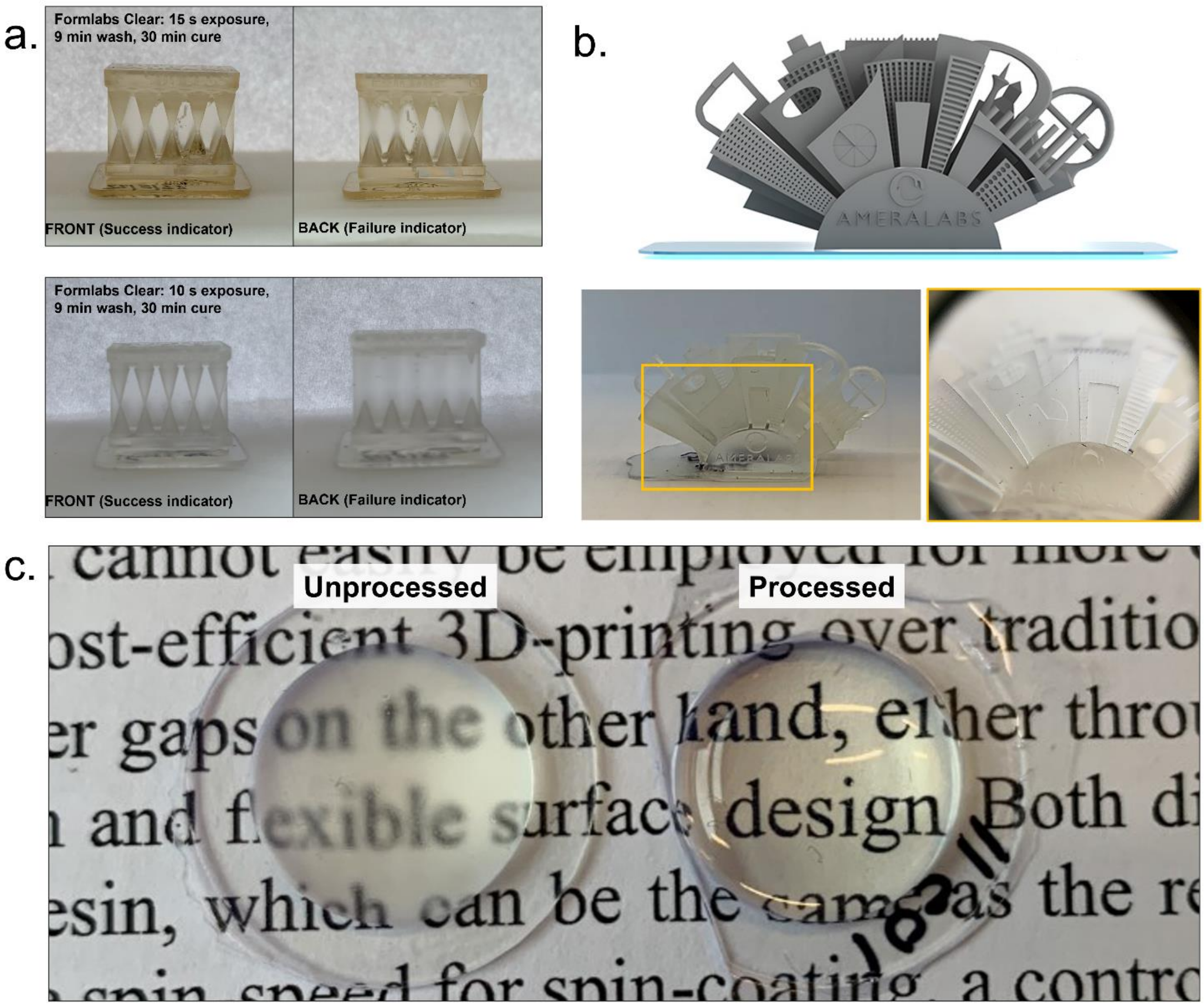


**Supplementary Figure 1. 3D Printer Calibration and Lens Printing. (a)** Exemplary prints using the TableFlip Foundry Cones of Calibration tool. Over-exposed print settings are identified by extra cones on the Fail indicator and yellowing of the print resin (TOP), and successful optimisation is determined by only one row of bottom cones on the Fail indicator (BOTTOM). **(b)** Secondary validation was achieved using the Ameralabs Town resolution test. The optimised settings resulted in the finest details being resolved on the Ameralabs print, down to 50 μm when resolved using a 10× magnifying hand lens. **(c)** Unprocessed (LEFT) and processed 3D printed lenses (RIGHT), demonstrating the increased optical quality following processing.

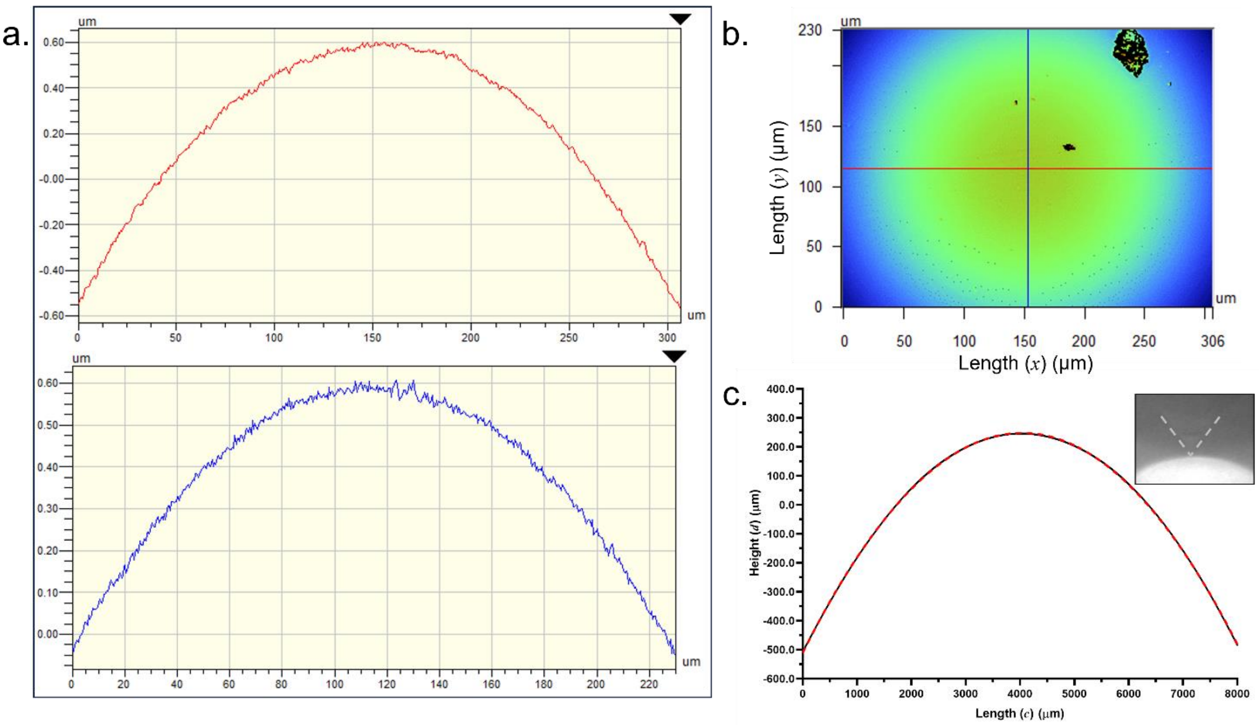


**Supplementary Figure 2. Surface Curvature Measurements using White Light Interferometry and Stylus Profilometry. (a)** Perpendicular measurements of surface curvature over a 300 μm field of view measured from **(b)** a white light interferometry image. **(c)** a stylus profilometry measurement of a 3D printed lens surface over an 8000 μm long trace (cut out shows stylus tip interacting with convex lens surface).

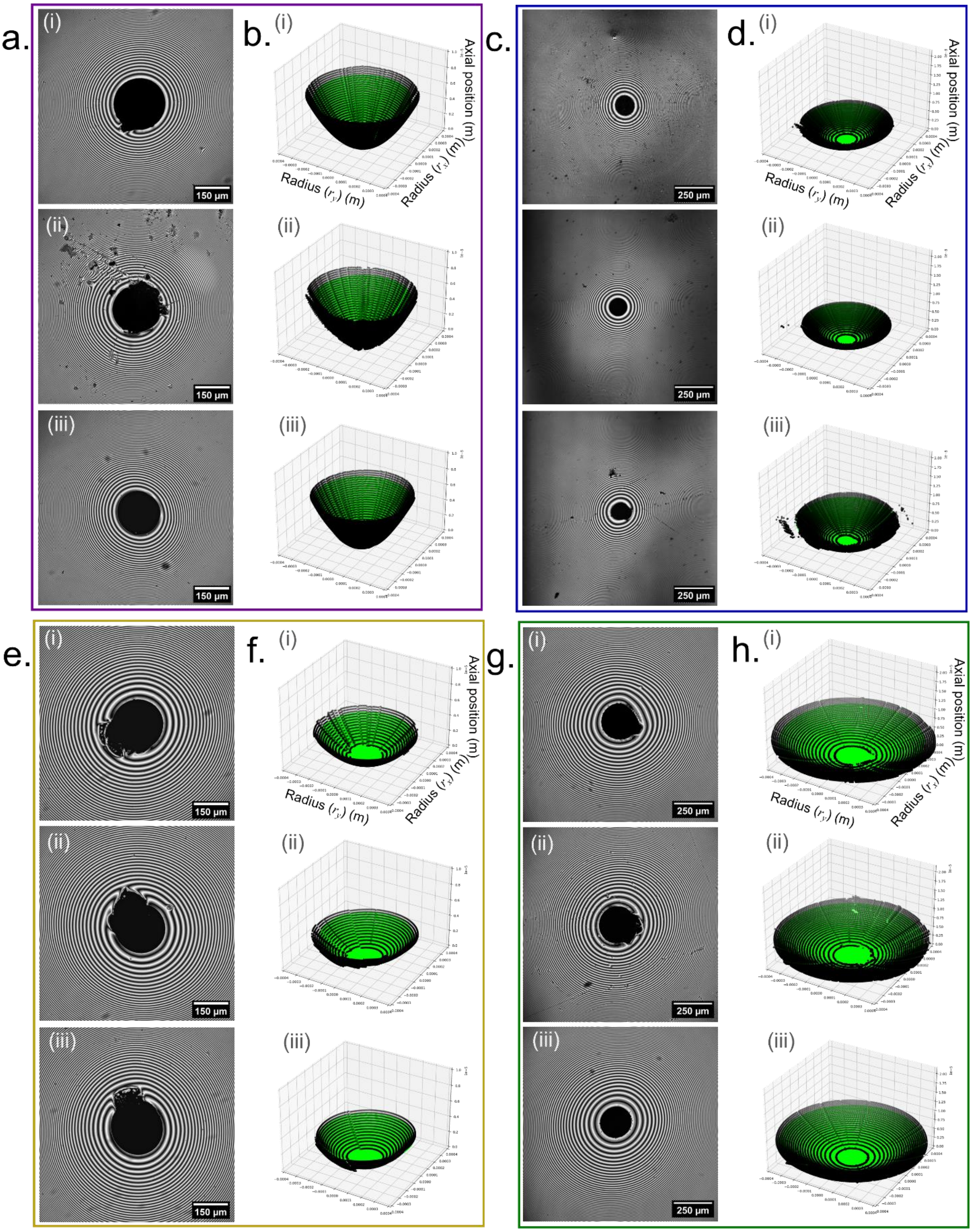


**Supplementary Figure 3. Surface Curvature Reconstructions using IRM Image Data.** The 2D IRM images and 3D surface reconstructions are presented for three replicate 3D printed lenses for various prescriptions: **(a)** $f = +\ 12.5$ mm, **(b)** $f = +\ 19.9$ mm, **(c)** $f = +\ 35.0$ mm, and **(d)** $f = +\ 49.8$ mm.

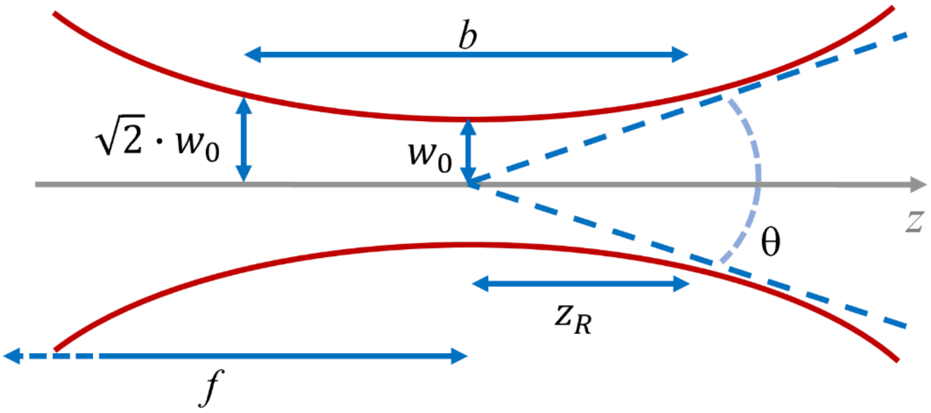


**Supplementary Figure 4. Gaussian Beam Metrics.** A schematic of Gaussian beam width ($w_z$) along the optical axis at the focal length ($f$), showing the beam waist ($w_0$), the Rayleigh range ($z_R$), the angular spread ($\theta$), and the depth of focus ($b$).

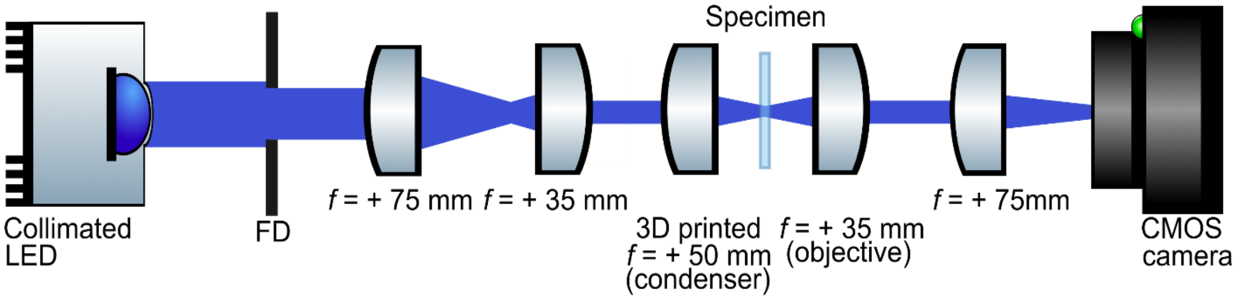


**Supplementary Figure 5. Brightfield Transmission Microscope Schematic.** A schematic of a brightfield transmission microscope setup used to demonstrate the imaging capabilities of 3D printed lenses (not presented to scale). An aperture stop acted as a field diaphragm (FD) controlling the diameter of a collimated blue LED source ($\lambda$ = 470 nm). A pair of $f$ = + 75 mm and $f$ = + 35 mm lenses reduced the beam size to fill the numerical aperture of the 3D printed condenser lens ($f$ = + 50 mm). Transmitted light was collected using an $f$ = + 35 mm objective lens and focussed onto a CMOS camera sensor using an $f$ = + 75 mm lens.
